# Specific host translation perturbations promote Tn*Smu*1 loss independently of the canonical ImmA regulator

**DOI:** 10.64898/2026.09.15.751695

**Authors:** Liang Bao, Todd Kitten, Ping Xu

## Abstract

Integrative and conjugative elements (ICEs) are major drivers of horizontal gene transfer and bacterial genome evolution. Although ICE-encoded regulatory circuits have been extensively characterized, the impact of host physiology on the stability of integrated ICEs remains poorly understood. Here, we identify a host-dependent pathway that links specific host translation perturbations to loss of the ICE Tn*Smu1* in *Streptococcus mutans*. Analysis of host-gene deletion mutants revealed that disruption of *fmt*, *rnjA*, or *rnjB*—three translation-associated host genes—reproducibly promoted Tn*Smu1* loss through a mechanism that bypasses the canonical ICE-encoded metalloprotease ImmA but remains dependent on the native attachment site *attR*. This phenotype was selective, as mutations affecting other essential cellular functions, including protein folding, tRNA modification, cell division, and fatty acid biosynthesis, failed to destabilize Tn*Smu1* despite undergoing the same experimental evolution and accumulating adaptive genomic changes. Preventing Tn*Smu1* loss in these translation-associated mutants markedly reduced bacterial growth, whereas loss of the element improved fitness, indicating that ICE elimination alleviates the cost associated with Tn*Smu1* retention under these conditions. Finally, we show that the relationship between host translation and Tn*Smu1* stability extends to a genetically distinct *S. mutans* clinical isolate, although with strain-dependent penetrance. Together, these findings identify host translational state as an important physiological determinant of Tn*Smu1* stability and reveal that bacterial hosts can influence the maintenance of integrated mobile genetic elements through mechanisms that extend beyond element-encoded regulatory circuits.

## Introduction

Horizontal gene transfer (HGT) is a major driver of bacterial evolution, enabling the rapid dissemination of antibiotic resistance, virulence determinants, and metabolic traits(1–3). HGT occurs through three major mechanisms—transformation, transduction, and conjugation—each relying on distinct molecular processes and genetic determinants(4). Among the mobile genetic elements that mediate HGT, integrative and conjugative elements (ICEs) are particularly widespread and play an important role in bacterial genome diversification(2, 5–8). ICEs reside stably integrated within the bacterial chromosome, where they replicate passively with the host genome. Upon activation, they excise from the chromosome, circularize, replicate in the donor cells, are transferred to recipient cells by conjugation, and integrate into the recipient genome, thereby completing their horizontal transmission cycle(2, 7).

The life cycle of ICEs is tightly regulated because it determines both the persistence of the element within its current host and its capacity for horizontal dissemination. Current models have focused primarily on regulatory circuits encoded by the ICE itself. For example, the ImmR/ImmA module controls excision of Tn*Smu1* and ICE*Bs1*(2, 9, 10), whereas Tn*916* employs an Orf9/antisense RNA regulatory system(7). However, host-encoded factors can also influence ICE dynamics. For example, RecA, together with ICE-encoded recombination proteins, promotes inter-ICE recombination and contributes to the formation of mosaic SXT/R391 ICE genomes(11). In addition, ICE excision and replication can engage the host SOS response, demonstrating that ICE activity can be coupled to broader host stress pathways(12). These findings indicate that ICE dynamics are shaped by both element-encoded regulatory circuits and host cellular processes. Whether host physiological pathways actively influence the stability and loss of integrated ICEs, however, remains poorly understood.

Translation is a central and highly regulated component of bacterial physiology (13–15). Changes in the translational machinery, ribosome function, and RNA metabolism can rapidly reshape global protein synthesis and cellular physiology, particularly during adaptation to environmental or metabolic stress(13). Because translation is tightly integrated with cellular growth and homeostasis, perturbation of translation-associated processes could provide a physiological interface through which the host influences the maintenance of integrated mobile genetic elements. However, whether host translational state affects ICE stability or loss remains unclear.

*Streptococcus mutans*, a primary etiologic agent of dental caries and an opportunistic cause of infective endocarditis, provides an excellent model for addressing this question(9, 10, 16). The oral microbiome is recognized as a hotspot for horizontal gene transfer, and *S. mutans* harbors multiple mobile genetic elements that contribute to genome diversification(9, 10, 16). Among these is the ICE Tn*Smu1*. Excision of Tn*Smu1* is regulated by the conserved ImmR/ImmA repressor system(9, 10). Whether host translation-associated processes also contribute to Tn*Smu1* stability has not been examined. Building on our previous success in generating viable mutants of genes previously classified as essential(17), we asked whether disruption of core cellular functions(18) affects Tn*Smu1* maintenance. Working in strain UA159, we found that disruption of three translation-associated host genes—*fmt*, *rnjA*, and *rnjB*—reproducibly promoted adaptive loss of Tn*Smu1*, whereas mutations affecting other essential cellular processes did not. We further show that this host-mediated response requires the *attR* recombination site but bypasses the canonical ImmA protease, and that preventing Tn*Smu1* loss under these translation-associated perturbations substantially reduces bacterial fitness. Finally, we demonstrate that the host translation–ICE stability relationship extends to a genetically distinct clinical isolate of *S. mutans*, although with strain-dependent penetrance. Together, these findings identify host translational state as an important physiological determinant of ICE stability and reveal that core host processes, in addition to ICE-encoded regulatory circuits, govern the maintenance of an integrated mobile genetic element.

## Results

### Specific host translation perturbations promote Tn*Smu1* loss

Tn*Smu1* is a ∼20-kb ICE comprising at least 30 predicted open reading frames in *S. mutans* UA159 (Figure 1A). Under the canonical regulatory model, Tn*Smu1* activation is controlled by the ImmR/ImmA module, in which the metalloprotease ImmA cleaves the ImmR repressor to initiate excision (9, 10, 16). Excision occurs through site-specific recombination between the left and right attachment sites (*attL* and *attR*)(19), which share a 17-bp direct repeat(9). The *attL* site lies within the 3′ end of the *tRNA-Leu* gene, whereas *attR* is located in an intergenic region (Figure 1A) (9).

**Figure 1.**
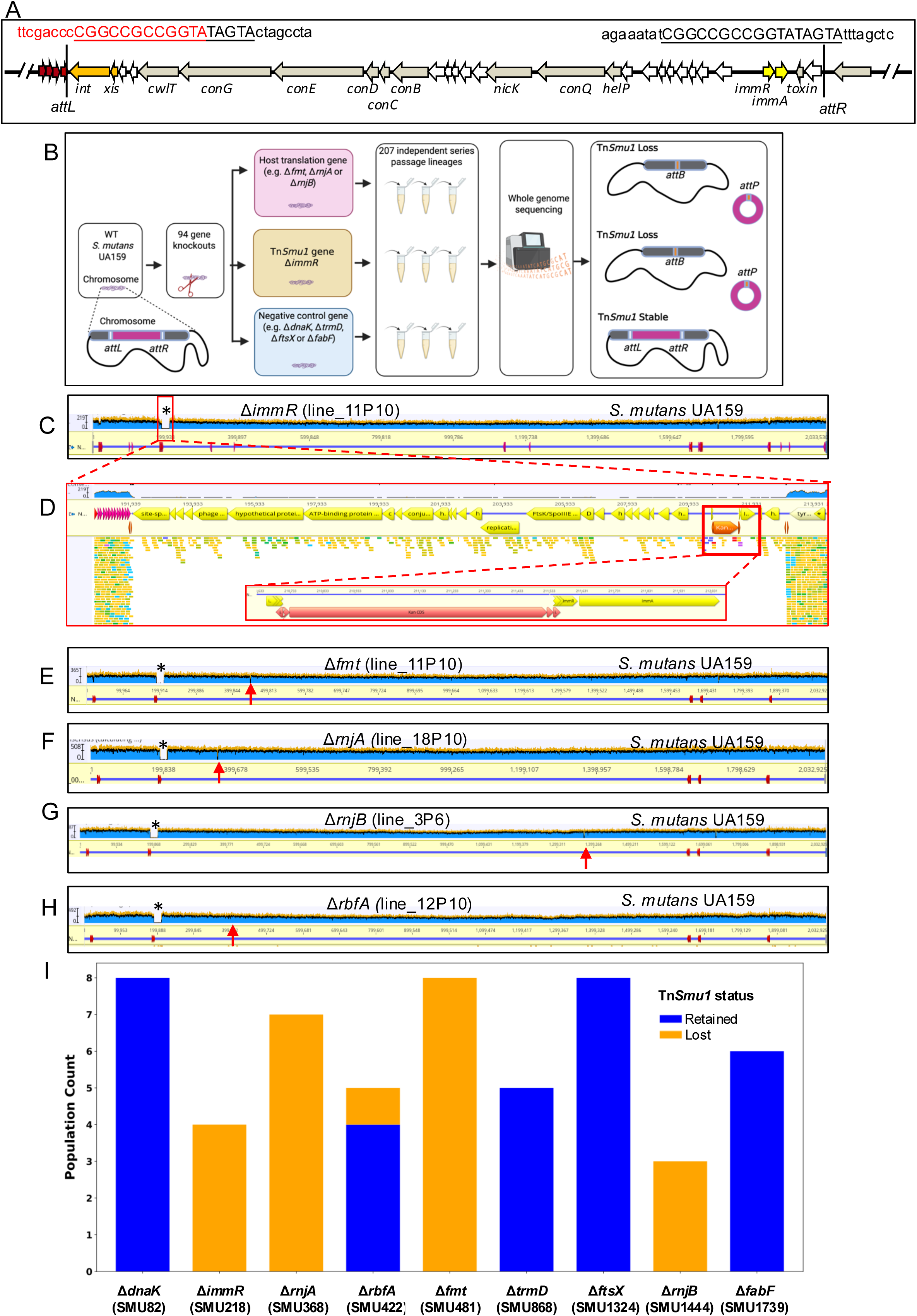
Analysis of host-gene deletion mutants identifies translation-associated host factors linked to Tn*Smu1* stability. **(A)** Genetic organization of the ICE Tn*Smu1* in *S. mutans* UA159. The 17-bp direct repeats corresponding to *attL* and *attR* are underlined and shown in uppercase letters, whereas the flanking chromosomal sequences are shown in lowercase. *attL* is located within the 3′ end of the tRNA-Leu gene, with the tRNA-derived nucleotides highlighted in red, whereas *attR* resides in an intergenic region. The integrase (*int*) and excisionase (*xis*) genes are shown in orange, and the regulatory genes *immR* and *immA* are shown in yellow. **(B)** Experimental workflow. Ninety-four targeted deletion mutants were constructed. For each mutant, 1–10 independent experimental evolution lineages were established, yielding a total of 207 evolved lineages. Evolved populations were analyzed by whole-genome sequencing. The image was created in BioRender. Bao, L. (2027) https://BioRender.com/61n2m6z **(C–H)** Representative whole-genome sequencing read coverage across the *S. mutans* UA159 chromosome after mapping to the reference genome (NC_004350). Regions marked by asterisks indicate loss of the ∼20-kb Tn*Smu1* element in the canonical control (Δ*immR*) and in mutants affecting translation-associated host processes (Δ*fmt*, Δ*rnjA*, Δ*rnjB*, and Δ*rbfA*). Red arrows indicate the locations of the deleted genes. The Δ*immR* lineage (C and D) was mapped to a modified reference genome in which *immR* was replaced by a kanamycin resistance cassette (coordinates 1–2,033,530 bp), whereas the Δ*fmt* (**E**), Δ*rnjA* (**F**), Δ*rnjB* (**G**), and Δ*rbfA* (**H**) lineages were mapped to the original NC_004350 reference genome (coordinates 1–2,032,925 bp). Vertical red arrow, site of original gene deletion; the height of the blue segments indicates the number of sequence reads mapped to the reference sequence at the coordinates shown on the X axis. **(I)** Summary of Tn*Smu1* retention and loss frequencies across independently evolved mutant lineages. Gene names are shown below each bar, with the corresponding *S. mutans* UA159 locus tags indicated in parentheses. Blue indicates Tn*Smu1* retention and orange indicates Tn*Smu1* loss. Tn*Smu1* loss was observed in 4/4 Δ*immR* (SMU218), 7/7 Δ*rnjA* (SMU368), 1/5 Δ*rbfA* (SMU422), 8/8 Δ*fmt* (SMU481), and 3/3 Δ*rnjB* (SMU1444) lineages, whereas Δ*dnaK* (SMU82; 0/8), Δ*trmD* (SMU868; 0/5), Δ*ftsX* (SMU1324; 0/8), and Δ*fabF* (SMU1739; 0/6) retained Tn*Smu1* in all lineages examined.

To identify host factors associated with ICE stability, we targeted 309 genes previously classified as essential in *S. mutans* UA159 and successfully constructed deletion mutants for 88 of them, together with six nonessential-gene deletion mutants, yielding 94 mutants in total (Figure 1B; Table S1)(18). Notably, many of these viable mutants, including Δ*fmt*, represent, to our knowledge, the first reported deletions in *S. mutans* of genes previously considered essential (Table S1)(18). For each mutant, one to ten independent lineages were established, generating a total of 207 experimental-evolution populations that were serially passaged and analyzed by whole-genome sequencing (Figure 1B; Table S1).

As expected, deletion of the ICE-encoded repressor gene, *immR,* resulted in Tn*Smu1* loss in four independently evolved lineages (Figure 1C–D; Table S1). Remarkably, apart from this positive control, reproducible Tn*Smu1* loss was confined to a subset of mutants affecting translation-associated host processes (Figure 1E–I; Table S1). Deletion of *fmt*, encoding methionyl-tRNA formyltransferase(20), resulted in Tn*Smu1* loss from all eight independently evolved lineages (Figure 1E and 1I; Table S1). Likewise, deletion of *rnjA* or *rnjB*, which encode the two RNase J paralogs involved in mRNA maturation (21, 22), resulted in Tn*Smu1* loss in all seven and three independently evolved lineages, respectively (Figure 1F, G, and I; Table S1). In contrast, deletion of *rbfA*, encoding a 30S ribosome maturation factor(23), produced only partial penetrance, with Tn*Smu1* loss occurring in one of five independently evolved populations (Figure 1H and I; Table S1).

To verify that Tn*Smu1* destabilization resulted specifically from disruption of *fmt*, we took advantage of a Δ*fmt* mutant that retained a wild-type copy of *fmt*. This strain, designated Δ*fmt*-*WTCR*, (for “wild type copy retained” (17)), arose spontaneously during construction of the Δ*fmt* mutant, when the chromosomal region containing *fmt* duplicated before allelic replacement of one copy of *fmt* with a kanamycin-resistance cassette. Thus, Δ*fmt*-*WTCR* retained one wild-type copy of *fmt* while otherwise undergoing the same allelic-replacement procedure and subsequent strain handling as the Δ*fmt* mutant. It therefore provided an internal genetic control for distinguishing effects caused by loss of *fmt* from those associated with strain construction or subsequent passage (Fig. S1A–B). Unlike the complete Δ*fmt* mutant, the Δ*fmt*-*WTCR* retained Tn*Smu1* throughout serial passage (Fig. S1A–B), supporting the conclusion that Tn*Smu1* destabilization was associated with loss of *fmt* rather than with the strain-construction or passage history alone. To exclude potential polar effects on neighboring genes, we analyzed four different constructions of deletion mutants. Both a Δ*fmt* mutant that also removed the 5′ portion of the downstream gene SMU482 (Fig. S1C–D; N=7, Table S1) and a newly generated Δ*fmt* mutant that left SMU482 intact (Fig. S1E–F; N=1, Table S1) exhibited Tn*Smu1* loss. In contrast, deletion of the downstream gene SMU482 alone (Fig. S1G–H; N=1, Table S1) or the upstream gene SMU480 alone (Fig. S1I–J; N=2, Table S1) did not destabilize Tn*Smu1*. Together, these results support a specific association between *fmt* disruption and Tn*Smu1* destabilization and argue against polar effects on adjacent genes as the underlying cause.

To determine whether Tn*Smu1* loss emerged progressively during serial passage, we examined Tn*Smu1* abundance, based on sequencing reads aligned to the reference genome, in three independent Δ*fmt* lineages (line_2, line_11, and line_14) at passage 1. Two of the three lineages, line_11 and line_14 (line_11 shown in Fig. S1C–D), exhibited >50% reductions in Tn*Smu1* coverage relative to the chromosomal background, whereas line_2 showed a <50% reduction (Fig. S1K–L). During subsequent passages, Tn*Smu1* coverage declined further, with >50% reductions observed by passage 10 in line_11 and line_14 (line_11 shown in Fig. 1E) and by passage 3 in line_2 (Fig. S1E–F). These results indicate that Tn*Smu1* loss emerged progressively during experimental evolution rather than being uniformly established at the outset of the experiment.

To determine whether Tn*Smu1* loss simply reflected prolonged serial passage or adaptive genome evolution, we analyzed mutants affecting several additional cellular processes as negative controls, including protein folding (Δ*dnaK*), tRNA methylation (Δ*trmD*), cell division (Δ*ftsX*), and fatty acid synthesis II (Δ*fabF*). Although these mutants underwent the same experimental evolution protocol and accumulated adaptive genomic changes (Fig. S2A–J), none exhibited Tn*Smu1* loss (0/8, 0/5, 0/8, and 0/6 evolved lineages, respectively; Figure 1I; Table S1). Whole-genome sequencing revealed multiple adaptive mutations, including recurrent chromosomal copy-number variations outside the Tn*Smu1* locus (Fig. S2A–J). For example, independently evolved Δ*trmD* lineages repeatedly acquired chromosomal amplifications ranging from 3.8 to 185.8 kb that encompassed *proS*, which encodes prolyl-tRNA synthetase (Fig S2C–J). Similar amplification of chromosomal regions encompassing *proS* has been implicated in adaptation to impaired tRNA methylation in *E. coli*(24). Likewise, in our previous study, mutants deleted for any of four genes in the aromatic amino acid synthase pathway consistently acquired chromosomal duplications encompassing a peptide transporter gene cluster (SMU255–SMU259)(25). Despite the obvious selective pressure that led to these adaptive genomic changes, Tn*Smu1* remained stably integrated in every lineage.

Collectively, these findings demonstrate that Tn*Smu1* destabilization is not a general consequence of serial passage or adaptive genome evolution. Instead, reproducible Tn*Smu1* loss is associated with specific perturbations of host translation-associated processes. The affected genes function in distinct processes, including tRNA formylation, RNA metabolism, and ribosome maturation, suggesting that Tn*Smu1* stability is sensitive to the translational state of the host rather than to disruption of a single molecular step. The absence of Tn*Smu1* loss in Δ*trmD* lineages further indicates that not all perturbations affecting translation-associated functions are sufficient to destabilize the element.

### Loss of kanamycin-marked Tn*Smu1* during serial passage of **Δ***immR* lineages

Because the Δ*immR* allele was marked with a kanamycin resistance cassette in our deletion construction, we independently verified Tn*Smu1* loss by monitoring the frequency of kanamycin-resistant cells during serial passage. By passages 3 and 7, the wild-type retained strain (Δ*immR*-*WTCR*), which carries a wild-type copy of *immR*, retained both Tn*Smu1* and kanamycin resistance (Figure 2A–B; Fig. S3A–D; Table S2). In contrast, all four independently evolved Δ*immR* lineages underwent rapid population turnover, with kanamycin-resistant cells declining to 2.8–17.4% by passage 3 and becoming nearly undetectable by passage 7 (below the limit of detection to 0.0008%; Figure 2A–B; Table S2). Whole-genome sequencing of kanamycin-sensitive colonies isolated from passage 7 confirmed loss of the ∼20-kb Tn*Smu1* locus (Fig. S3E–H, N=4). The stable retention of both Tn*Smu1* and kanamycin resistance in the Δ*immR*-*WTCR* strain supports the conclusion that loss of *immR* promotes Tn*Smu1* excision and destabilization. Together, these findings indicate that disruption of ImmR-mediated repression imposes a substantial fitness burden, resulting in strong positive selection for Tn*Smu1* loss.

**Figure 2.**
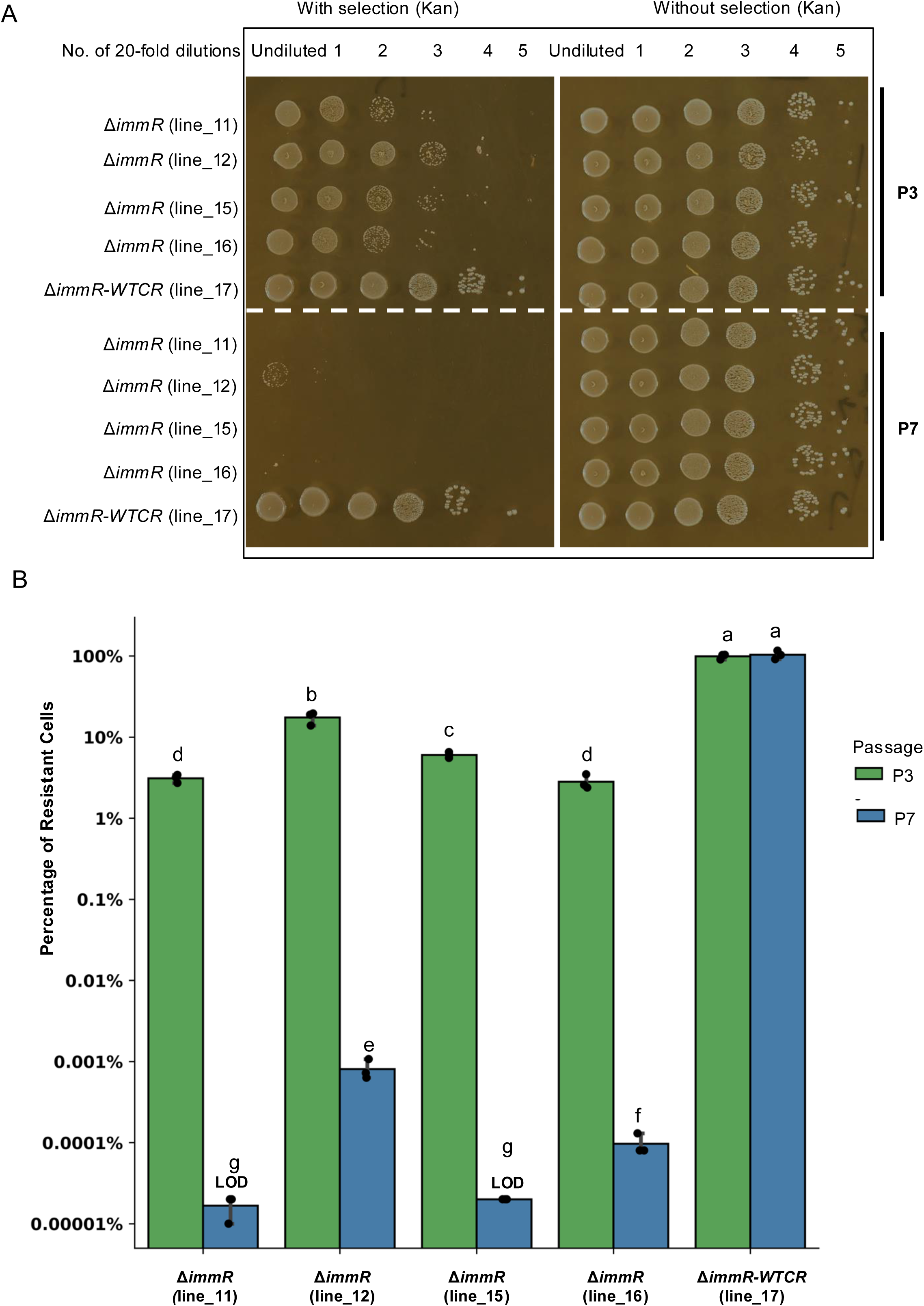
*immR* deletion imposes strong selection for Tn*Smu1* loss. **(A)** Growth of Δ*immR::kan* and the Δ*immR::kan-WTCR* strain on selective and nonselective media. At passage 3 and 7, cultures from four independently evolved Δ*immR* lineages (_11, _12, _15 and _16) and one Δ*immR-WTCR* lineage (_17) were serially diluted. Two microliters of each dilution were spotted onto BHI agar with or without kanamycin and incubated anaerobically at 37°C for 48 h. Dilution factors were indicated in the figure. **(B)** Quantification of the proportion of kanamycin-resistant and kanamycin-sensitive cells in the evolved Δ*immR* and Δ*immR-WTCR* lineages at passage 3 and 7. Values represent the mean ± SD from three biological replicates. Statistical significance was determined using one-way ANOVA with Tukey’s HSD test; groups sharing a common letter are not significantly different (*P* > 0.05). Note that the CFU for two Δ*immR* lineages (_11 and _15) at P7 was set at the limit of detection (LOD) since no colonies were recovered by assuming that one colony formed in the most concentrated sample that was plated.

### Tn*Smu1* loss following *immR* deletion requires *attR*-dependent recombination

Because the Δ*immR* populations became heterogeneous for Tn*Smu1* retention during serial passage, and because Δ*immR* cells exhibited smaller colonies than the Δ*immR*-*WTCR* strain on kanamycin-containing plates (Figure 2A) and reduced fitness relative to ΔTn*Smu1* cells (Figure 2B), we next quantified the growth of the Δ*immR* populations under kanamycin selection. Passage 2 lineages were cultured overnight in BHI containing kanamycin to maintain selection for Δ*immR* or Δ*immR*-*WTCR*, then diluted into fresh, selective BHI. Growth monitoring over 12 hours revealed that the Δ*immR* lineage grew significantly slower than the Δ*immR*-*WTCR* control (Figure 3A; Table S3), confirming that the disruption of *immR* causes a distinct fitness defect.

**Figure 3.**
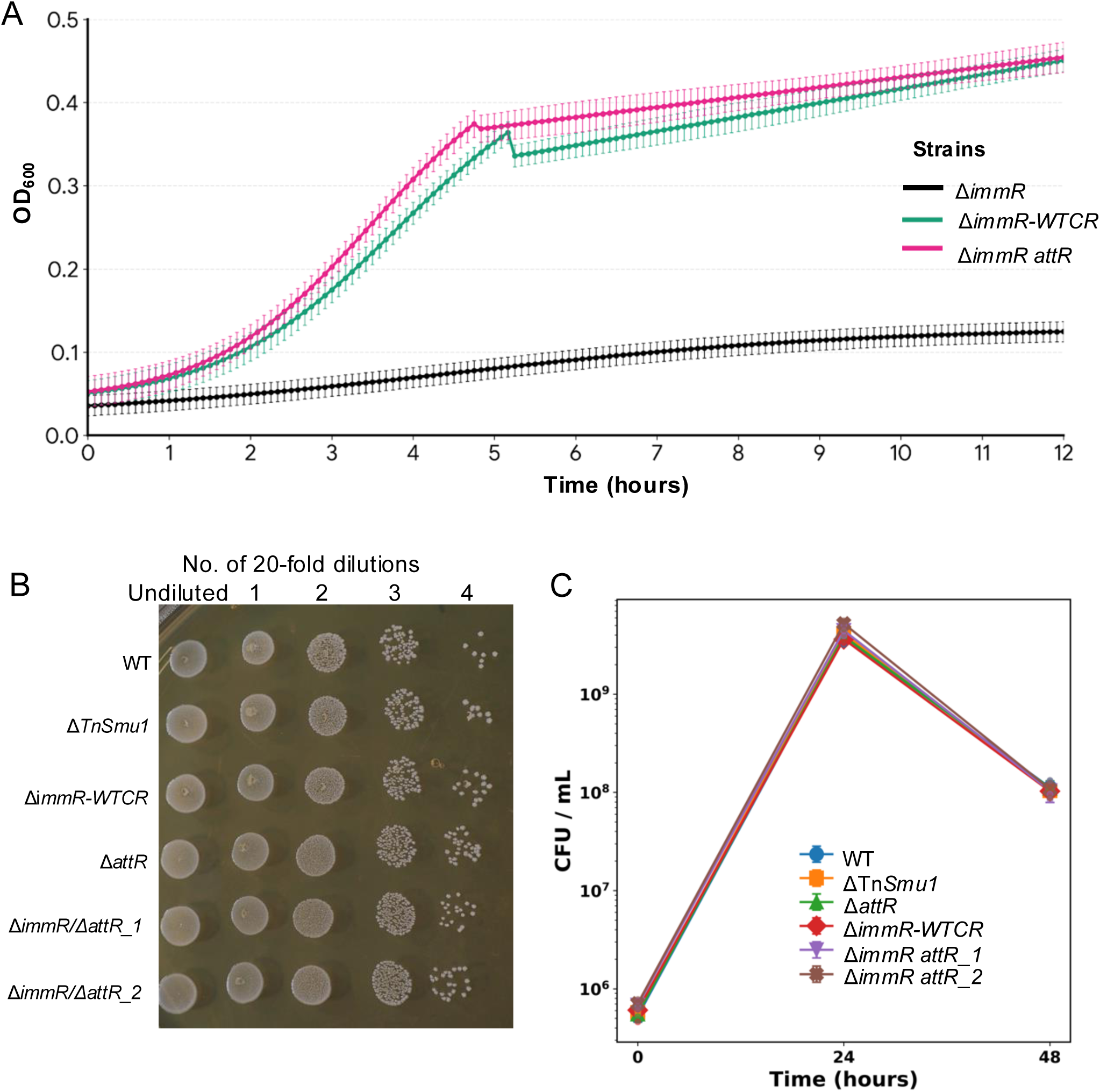
Blocking Tn*Smu1* loss through *attR* deletion suppresses the growth defect associated with *immR* deletion. **(A)** Growth kinetics of Δ*immR* (line_15), Δ*immR*-WTCR (line_17), and Δ*immR* Δ*attR* (line_11) strains. Passage-derived cultures were grown in BHI supplemented with kanamycin, and growth was monitored for 12 h by measuring OD₆₀₀ with a microplate reader. Solid lines represent the mean optical density of triplicate wells (n = 3), and shaded areas indicate the standard deviation (± SD). **(B)** Spot-dilution analysis of strains retaining or lacking Tn*Smu1*. Twenty-four-hour cultures were serially diluted, and 2 μL of each dilution was spotted onto BHI agar without antibiotics and incubated anaerobically at 37°C for 48 h. The strains tested were wild-type *S. mutans* UA159 (WT), a Tn*Smu1*-free derivative isolated from the Δ*immR* lineage (ΔTn*Smu1*; line_11P7_kanfree; Fig. S3E–F), Δ*attR*, Δ*immR*-WTCR (line_17), and two independently passaged Δ*immR* Δ*attR* lineages (line_11 and line_12). Representative whole-genome sequencing profiles of the Δ*immR*-WTCR and Δ*immR* Δ*attR* lineages are shown in Fig. S3A–D and Fig. S3I–L, respectively. **(C)** Growth curves of WT, ΔTn*Smu1*, Δ*attR*, Δ*immR-WTCR*, and two lineages of Δ*immR* Δ*attR*. Viable cell numbers were determined by CFU enumeration at 0, 24, and 48 h. Data represent the mean ± SD from three biological replicates.

To determine whether Tn*Smu1* loss following *immR* deletion requires *attR*-dependent recombination, we replaced the *attR* attachment site with an erythromycin-resistance cassette before introducing the Δ*immR* mutation, generating the Δ*immR* Δ*attR* strain. Whole-genome sequencing of two independently passaged lineages confirmed retention of Tn*Smu1* in the Δ*immR* Δ*attR* background, demonstrating that deletion of *attR* prevents the loss of the integrated ICE that occurs following *immR* deletion (Fig. S3I–L). Thus, the canonical ImmR regulatory pathway requires the *attR* recombination site to eliminate the integrated ICE.

Unexpectedly, preventing Tn*Smu1* excision also eliminated the severe growth defect associated with Δ*immR*. The Δ*immR* Δ*attR* strain exhibited growth comparable to that of the wild type under standard laboratory conditions (Figure 3A–C; Table S3–S4). These findings place *immR* genetically upstream of *attR*-mediated recombination and indicate that the pronounced fitness defect caused by loss of *immR* depends on successful ICE excision rather than on *immR* deletion alone.

### Adaptive Tn*Smu1* loss improves bacterial fitness under translation-associated perturbations

To determine whether Tn*Smu1* loss provides a selective advantage under host translation perturbation, we genetically prevented ICE excision by deleting the *attR* site. In this excision-deficient background (Δ*fmt* Δ*attR*), Tn*Smu1* remained stably integrated throughout serial passage (Fig. S4A; Fig. S5A–B). We then assessed bacterial fitness by measuring viable cell counts (CFU mL⁻¹) at 0, 24, and 48 h (Figure 4 A–B).

**Figure 4.**
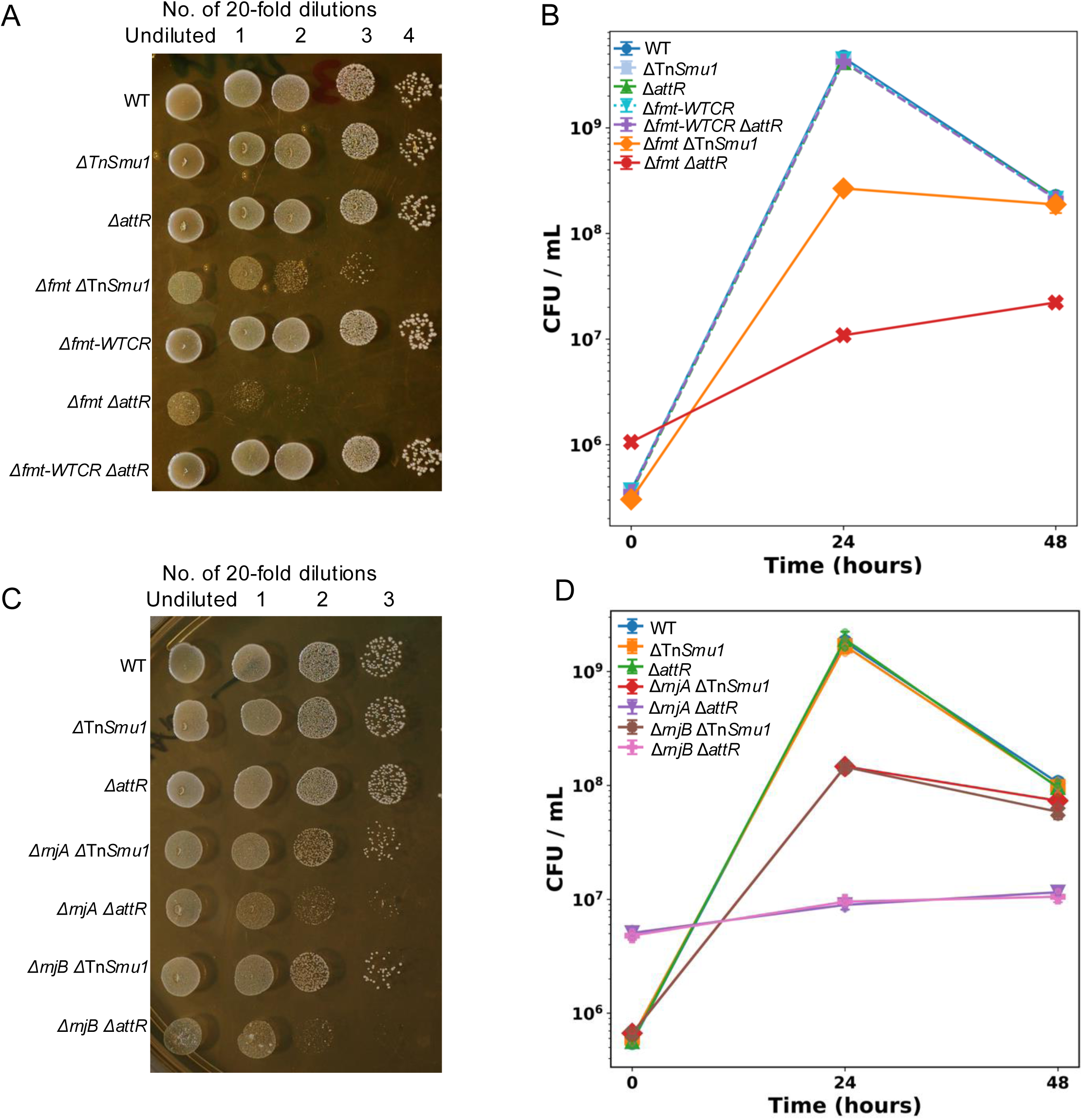
Retention of Tn*Smu1* imposes a substantial fitness cost in translation-associated mutant backgrounds. **(A)** Cultures of the strains indicated grown for 24 h were serially diluted, and 2 μL of each dilution were spotted onto BHI agar and incubated anaerobically at 37°C for 48 h. Dilution factors are indicated in the figure. Additional details are provided in the Materials and Methods. **(B)** Growth curves of WT, ΔTn*Smu1*, Δ*attR*, Δ*fmt* ΔTn*Smu1*, Δ*fmt*-*WTCR*, Δ*fmt* Δ*attR*, and Δ*fmt*-*WTCR* Δ*attR*. Viable cell numbers were determined by CFU enumeration at 0, 24, and 48 h. Data represent the mean ± SD of three biological replicates. **(C)** Cultures of the strains indicated grown for 24 h were serially diluted, and 2 μL of each dilution were spotted onto BHI agar and incubated anaerobically at 37°C for 48 h. Dilution factors are indicated in the figure. **(D)** Growth curves of WT, ΔTn*Smu1*, Δ*attR*, Δ*rnjA* ΔTn*Smu1*, Δ*rnjA* Δ*attR*, Δ*rnjB* ΔTn*Smu1*, and Δ*rnjB* Δ*attR*. Viable cell numbers were determined by CFU enumeration at 0, 24, and 48 h. Data represent the mean ± SD of three biological replicates.

The Δ*fmt* Δ*attR* strain, in which the host translation defect persisted but Tn*Smu1* could no longer be eliminated, exhibited the most severe growth defect and the lowest final cell density of all strains examined (Figure 4A–B; Table S5). In contrast, independently evolved Δ*fmt* lineages that had naturally lost Tn*Smu1* (Figure 1I) showed substantially improved growth (Figure 4A–B; Table S5), indicating that elimination of the ICE partially rescued the fitness defect associated with loss of *fmt*. As controls, the Tn*Smu1*-free strain (ΔTn*Smu1*; derived from the Δ*immR* mutant) (Fig. S3E–F), the Δ*attR* single mutant, the Δ*fmt*-*WTCR* strain (Fig. S1A–B), and the Δ*fmt*-*WTCR* Δ*attR* strain (Fig. S4C–D) all exhibited growth comparable to that of the wild-type strain (Figure 4A–B; Table S5).

To determine whether this fitness advantage extended to other translation-associated mutants, we performed the same analysis in Δ*rnjA* and Δ*rnjB* backgrounds. Similar to the Δ*fmt* mutant, the excision-deficient strains Δ*rnjA* Δ*attR* and Δ*rnjB* Δ*attR* exhibited substantially greater growth defects and lower final cell densities than the corresponding passaged Δ*rnjA* ΔTn*Smu1* and Δ*rnjB* ΔTn*Smu1* strains, which had spontaneously eliminated Tn*Smu1* during experimental evolution (Figure 4C–D; Table S6).

Collectively, these findings demonstrate that Tn*Smu1* retention imposes a substantial fitness burden in these translation-associated mutant backgrounds, whereas adaptive loss of the ICE alleviates this burden and provides a clear selective advantage to the host.

### Host-mediated Tn*Smu1* loss bypasses ImmA but requires *attR*-dependent recombination

Because the growth analyses in Figures 3 and 4 indicated that *attR* is required for both canonical ImmR derepression and host-mediated Tn*Smu1* loss, we next asked whether these phenotypes were reproducible across independently evolved lineages. Whole-genome sequencing of two to three independently passaged Δ*attR* lineages confirmed retention of Tn*Smu1* in Δ*immR* Δ*attR*, Δ*fmt* Δ*attR*, Δ*rnjA* Δ*attR*, and Δ*rnjB* Δ*attR* strains (Figure 5A; Fig. S3I–L; Fig. S5A–F; Table S7). These results independently validate that Tn*Smu1* loss driven by either ImmR disruption or specific host translation-associated perturbations requires the native *attR* site, consistent with site-specific recombination.

**Figure 5.**
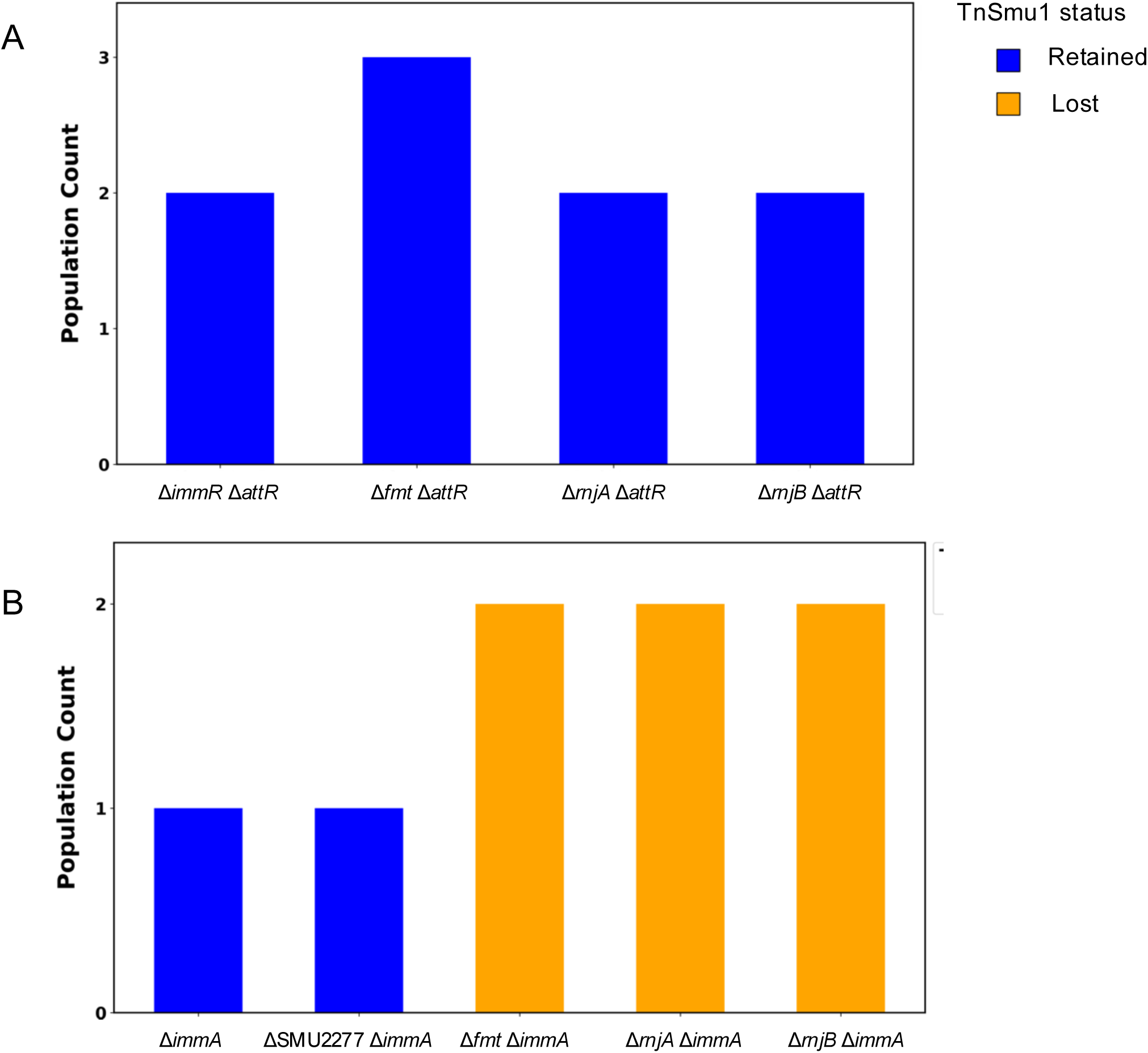
Host-mediated Tn*Smu1* loss requires *attR* but bypasses the canonical ImmA pathway. Whole-genome sequencing was performed on all evolved strains. **(A)** Tn*Smu1* status in the Δ*attR* background following serial passage. **(B)** Tn*Smu1* status in the Δ*immA* background following serial passage.

We next examined whether host-mediated Tn*Smu1* loss requires the canonical ImmA metalloprotease. As expected, deletion of *immA* alone did not trigger Tn*Smu1* loss during serial passage. As a control, we examined a ΔSMU277 Δ*immA* strain because disruption of *SMU277* did not cause Tn*Smu1* loss in our initial analysis (Table S1). SMU27*7* encodes a hypothetical protein, and its deletion provided a control for the effects of introducing a second host-gene mutation into the Δ*immA* background. ΔSMU277 Δ*immA* likewise retained the integrated Tn*Smu1*. In contrast, introduction of Δ*fmt*, Δ*rnjA*, or Δ*rnjB* into the Δ*immA* background still resulted in reproducible Tn*Smu1* elimination following experimental evolution (Figure 5B; Fig. S5G–P; Table S8).

Collectively, these findings confirm across multiple independently evolved lineages that Tn*Smu1* loss associated with specific host translation perturbations requires the native *attR* site but bypasses the canonical ImmA pathway. Together with the growth analyses in Figures 3 and 4, these results support a model in which specific host translation perturbations promote Tn*Smu1* loss through an ImmA-independent but *attR*-dependent mechanism (Figure 6).

**Figure 6.**
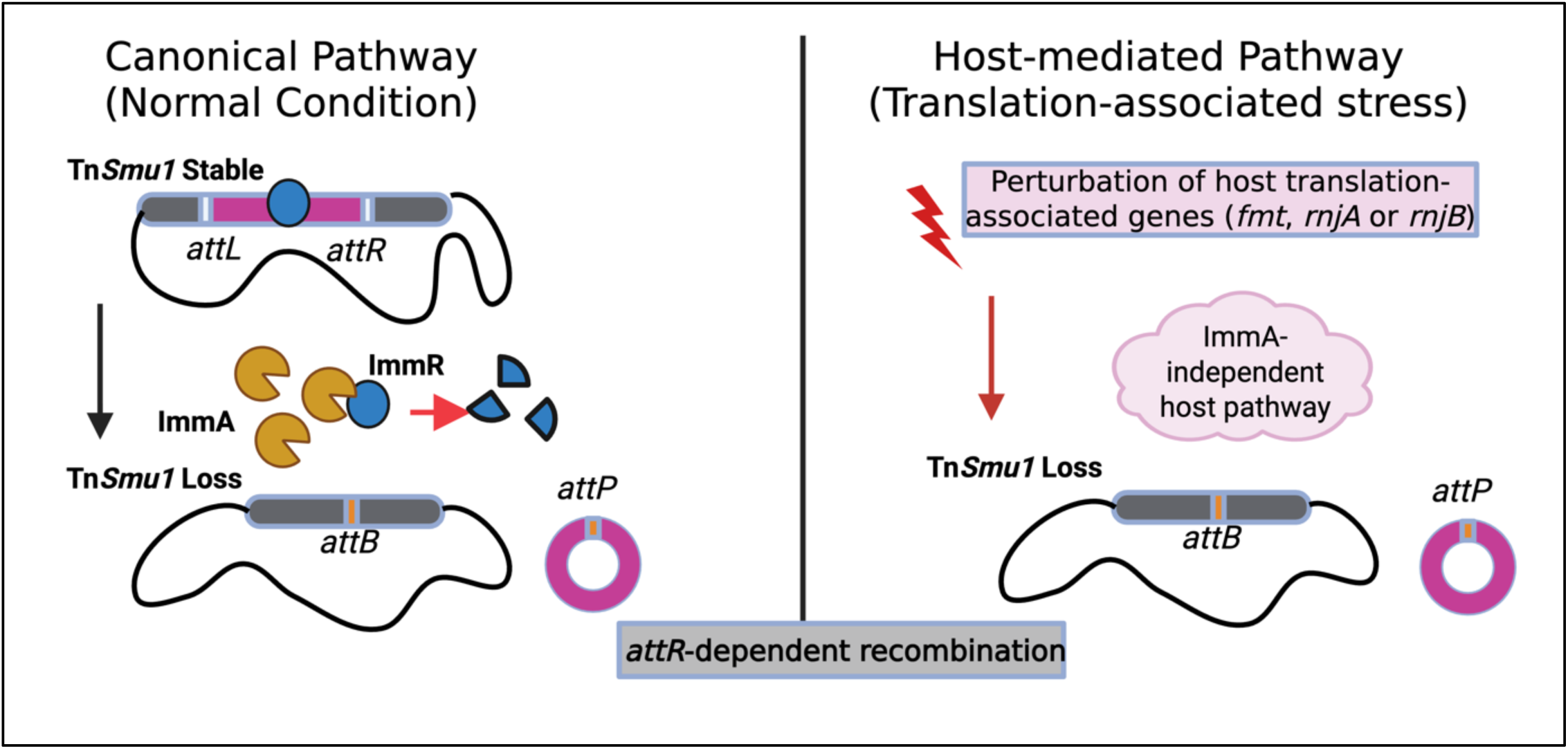
Model for host-mediated destabilization of integrated Tn*Smu1*. Under the canonical pathway, the Tn*Smu1*-encoded repressor ImmR maintains the element in an integrated state, whereas ImmA-mediated proteolysis of ImmR relieves repression and promotes excision (left). In contrast, disruption of translation-associated host genes (*fmt*, *rnjA*, and *rnjB*) promotes loss of chromosomally integrated Tn*Smu1* through a pathway that does not require ImmA. This host-mediated pathway may involve reduced ImmR synthesis or an alternative mechanism that destabilizes ImmR, although the molecular mechanism remains to be determined. Tn*Smu1* loss remains dependent on the native *attR* site, consistent with a requirement for site-specific excision. The subsequent fate of the excised Tn*Smu1* molecule was not determined in this study.

### Host translation influences Tn*Smu1* stability across genetically distinct *S. mutans* strains

To determine whether the relationship between host translation and Tn*Smu1* stability extends beyond the laboratory strain UA159, we examined *fmt*, *rnjA*, *rnjB*, and *rbfA* mutants in the genetically distinct bloodstream isolate *S. mutans* V403, which harbors an endogenous plasmid and causes both caries and endocarditis(27). Deletion of *fmt* or *rnjB* resulted in Tn*Smu1* loss in all independently evolved V403 lineages (3/3 and 4/4, respectively). Deletion of *rnjA* produced a less penetrant phenotype, with Tn*Smu1* loss detected in 2 of 5 lineages after extended evolution, whereas no Tn*Smu1* loss was observed in any of the three *rbfA* lineages examined (0/3; Fig. S6A–E; Table S9). Thus, disruption of *fmt* and *rnjB* reproducibly promotes Tn*Smu1* loss in two genetically distinct *S. mutans* backgrounds, while the effects of *rnjA* and *rbfA* are more strain- and context-dependent. The delayed and partial phenotype of Δ*rnjA* in V403, together with the absence of Tn*Smu1* loss following *rbfA* deletion in V403 despite its partial phenotype in UA159, indicates that the host translation–ICE stability relationship extends across strains but exhibits strain-dependent differences in penetrance. These differences may reflect variation in the genetic or physiological responses to specific translation-associated perturbations between the two strains.

## Discussion

Integrative and conjugative elements have traditionally been viewed as self-regulated mobile genetic elements whose maintenance and dissemination are governed primarily by element-encoded regulatory circuits (5, 6). In this study, we demonstrate that the bacterial host also plays an active role in determining ICE stability. By analyzing a broad collection of host-gene deletion mutants during experimental evolution, we identified host translational state as an important physiological determinant of the maintenance of the endogenous ICE Tn*Smu1* in *S. mutans* (Figure 1A–B). Disruption of three translation-associated host genes, *fmt*, *rnjA*, and *rnjB*, reproducibly promoted adaptive Tn*Smu1* loss, revealing an additional layer of host control over integrated mobile genetic elements.

A notable feature of this response is its specificity. Although mutations in *fmt*, *rnjA*, and *rnjB* perturb translation-associated processes through distinct molecular mechanisms, they all destabilized Tn*Smu1* (Figure 1C–I). In contrast, control mutants affecting protein folding, tRNA modification, cell division, and fatty acid biosynthesis failed to induce ICE loss despite undergoing the same experimental evolution and accumulating adaptive genomic changes (Fig. S2A–L). These findings argue against a nonspecific stress response and instead suggest that Tn*Smu1* responds, directly or indirectly, to physiological changes associated with specific perturbations of host translation. Whether this response reflects altered tRNA formylation, ribosome assembly, RNA metabolism, translation efficiency, or another consequence of altered host translational state remains to be determined.

Our genetic analyses further demonstrate that host-mediated Tn*Smu1* loss occurs through a mechanism distinct from the canonical regulatory pathway. In previously described models, activation of Tn*Smu1* depends on ImmA-mediated cleavage of the ImmR repressor. In contrast, deletion of *fmt*, *rnjA*, or *rnjB* resulted in Tn*Smu1* loss even in the absence of ImmA, demonstrating that the canonical metalloprotease is dispensable for host-mediated Tn*Smu1* loss (Figure 5B). Nevertheless, deletion of the *attR* recombination site completely abolished Tn*Smu1* loss in all three translation-associated mutants tested, indicating that host-mediated loss requires the native *attR* site and is consistent with site-specific recombination rather than nonspecific chromosomal instability or direct deletion of the ICE locus (Figure 5A). Together, these findings support a model in which specific host translation-associated perturbations engage an alternative host-dependent route that bypasses the canonical ImmA regulator while remaining dependent on *attR*-mediated recombination (Figure 6).

Interestingly, preventing excision by deleting *attR* produced strikingly different phenotypic outcomes in the two conditions. In the canonical pathway, the Δ*immR* Δ*attR* mutant retained Tn*Smu1* yet exhibited nearly wild-type growth, indicating that constitutive derepression of the ICE alone imposes little measurable fitness cost when excision is blocked (Figure 3). In contrast, the Δ*fmt* Δ*attR,* Δ*rnjA* Δ*attR* and Δ*rnjB* Δ*attR* mutants displayed the most severe growth defects. These defects were substantially alleviated in the corresponding evolved Δ*fmt*, Δ*rnjA* or Δ*rnjB* populations that had eliminated Tn*Smu1*, which showed markedly improved fitness (Figure 4). These contrasting phenotypes indicate that the physiological consequences of ICE retention depend on the physiological context in which Tn*Smu1* excision occurs. Rather than simply derepressing ICE gene expression, these translation-associated perturbations appear to create a host physiological state in which maintaining integrated Tn*Smu1* becomes particularly costly. Thus, Tn*Smu1* loss under these conditions provides a fitness benefit by relieving this physiological burden, whereas ImmR derepression in the canonical pathway primarily promotes the normal ICE excision cycle.

The molecular basis of this host-dependent regulatory pathway remains to be elucidated. Altered host translational state could influence the abundance, stability, or activity of ICE regulatory components independently of the canonical ImmA pathway. Changes in protein synthesis or RNA metabolism could affect ImmR abundance or alter the production of the recombination proteins Int and Xis. Alternatively, broader physiological changes associated with these translation-associated perturbations could modify the cellular environment required for efficient ICE recombination. Distinguishing among these possibilities will require direct measurements of ICE transcript abundance, regulatory-protein levels, and recombination activity.

Our results further indicate that the relationship between host translation and Tn*Smu1* stability extends across genetically distinct *S. mutans* backgrounds but varies in penetrance. Deletion of *fmt* or *rnjB* reproducibly promoted Tn*Smu1* loss in both the laboratory strain UA159 and the bloodstream isolate V403 (Figure 1; Fig. S6), demonstrating that this host–ICE relationship is not restricted to a single genetic background. In contrast, the effects of *rnjA* and *rbfA* were more context dependent. In V403, Δ*rnjA* produced delayed and incompletely penetrant Tn*Smu1* loss (Table S9), whereas Δ*rbfA* failed to promote detectable element loss despite exhibiting a partially penetrant phenotype in UA159. These differences suggest that the effects of specific translation-associated perturbations on ICE stability are influenced by the broader genetic and physiological background of the host. Variation in compensatory pathways, RNA metabolism, ribosome biogenesis, or other strain-specific regulatory features could alter the magnitude or timing of this response. Thus, although the link between host translation and Tn*Smu1* stability is maintained across distinct *S. mutans* backgrounds, its penetrance is modulated by host context, further emphasizing that ICE maintenance reflects an interaction between element-encoded functions and host physiology.

More broadly, because the core translation machinery is highly conserved across bacteria and ImmR/ImmA-like regulatory components occur in diverse mobile genetic elements in Gram-positive bacteria, including ICE*Bs1* in *Bacillus subtilis*(28), *Streptococcus thermophilus* (ICE*St1* and ICE*St3*)(29), and *Staphylococcus aureus* (Sa*PI3*)(30), related host-dependent mechanisms may contribute to the stability and maintenance of mobile genetic elements in other bacterial species. Whether similar perturbations of host translation influence these elements remains to be determined, but our findings raise the broader possibility that core host physiology constitutes an additional layer of mobile-element regulation.

In conclusion, our findings identify host translational state as an important physiological determinant of Tn*Smu1* stability and demonstrate that bacterial physiology contributes directly to the maintenance of integrated mobile genetic elements. Rather than acting solely as passive carriers of ICEs, bacterial hosts actively influence the balance between ICE maintenance and elimination through core physiological pathways. These results expand current models of ICE regulation and establish a new framework for understanding how host physiology shapes horizontal gene transfer and bacterial genome evolution. Elucidating this host-dependent regulatory pathway may ultimately provide new strategies to limit horizontal gene transfer and reduce the spread of bacterial virulence and antimicrobial resistance(31).

## Materials and Methods

### Wild-type parental strains and growth conditions

In this study, we primarily utilized *Streptococcus mutans* strain UA159, obtained from Ann Progulske-Fox (University of Florida). *S. mutans* strain V403 was provided by R. Facklam, Centers for Disease Control and Prevention and has been maintained as a frozen stock(32). *S. mutans* strains created in this study are described in the text. All bacterial cultures were routinely cultured in a BHI medium at 37°C under anaerobic conditions (0.2% O_2_, 9.9% H_2_, 9.9% CO_2_, and 80% N_2_), using a Coy anaerobic chamber, or stored at −80 °C for long-term storage.

### Transformation and mutant isolation

To prepare competent cells, *S. mutans* was inoculated into 1 mL of Todd–Hewitt broth supplemented with horse serum (TH+HS) and incubated anaerobically at 37°C overnight. The following day, 100 μL of the overnight culture was transferred into 1 mL of fresh TH+HS medium and incubated for an additional 3 h. Subsequently, 20 μL of this culture was inoculated into 1 mL of fresh TH+HS medium and incubated for 15 min to induce competence prior to the addition of transforming DNA.

Gene deletion constructs were generated by overlap extension PCR(33) using approximately 1-kb upstream and downstream homologous regions flanking the target coding sequence (CDS) together with either a kanamycin (*kan*) or erythromycin (*erm*) resistance cassette (primer sequences are listed in Table S10). For transformation, 50–500 ng of purified PCR product was mixed with 200 ng of competence-stimulating peptide (CSP; SGSLSTFFRLFNRSFTQALGK) and 150 μL of competent cells in either 96-well plates or 1.5-mL microcentrifuge tubes. Transformation reactions were incubated anaerobically at 37°C for 24 h. Following incubation, 2–5 μL of each transformation mixture was spotted onto BHI agar containing the appropriate selective antibiotic (kanamycin, 500 μg/mL; erythromycin, 10 μg/mL). After the inoculum had dried, plates were overlaid with 1 mL of molten BHI agar containing the corresponding antibiotic and allowed to solidify. Plates were incubated anaerobically at 37°C for 4–5 days to allow transformant colonies to develop.

Transformants were confirmed by PCR and, when indicated, by whole-genome sequencing. Genomic DNA was isolated from 2 mL of saturated cultures. Cells were harvested by centrifugation at 9,000 × *g* for 10 min at room temperature and resuspended in 200 μL of resuspension buffer (20 mM EDTA, 200 mM Tris-HCl, and 2% Triton X-100). Cell lysis was performed by adding 200 μL of Buffer AL (Qiagen, Cat. No. 19075). DNA was precipitated with 1 mL of 100% ethanol containing 100 mM sodium acetate, washed, air-dried, and resuspended in 100 μL of nuclease-free water for PCR analysis or whole-genome sequencing.

### Strain passage

Antibiotic-resistant transformants were inoculated into 1 mL of BHI supplemented with the appropriate antibiotic and incubated anaerobically at 37°C for 2–8 days until the culture reached an optical density at 600 nm (OD_600_) of 0.1–0.5. This initial culture was designated passage 0 (P0). A 300-μL aliquot of the P0 culture was then transferred into 3 mL of fresh BHI containing the same antibiotic and grown to saturation to generate passage 1 (P1).

To initiate subsequent passages, the P1 culture was mixed thoroughly by pipetting five times with a P1000 pipette, and 50 μL was transferred into 1 mL of fresh BHI containing the appropriate antibiotic. Cultures were incubated anaerobically at 37°C until reaching an OD_600_ 0.1 to 0.5, at which point the culture was designated as the next passage. This serial passaging procedure was repeated for the indicated number of passages, with each passage maintained in 1 mL of medium. For lineages requiring a larger volume at the final passage, 50 μL of the preceding passage was transferred into 3 mL of fresh BHI containing the appropriate antibiotic and grown to saturation. Passage cultures were preserved at −80°C in BHI containing 20% glycerol. Genomic DNA was isolated from the indicated passages for PCR analysis and/or whole-genome sequencing.

### Kanamycin resistance assay of Δ*immR* mutant strains

Δ*immR* or Δ*immR-WTCR* mutant strains were serially passaged in antibiotic-free BHI to monitor spontaneous loss of the kanamycin-marked Tn*Smu1* element. Briefly, passage 1 (P1) cultures were inoculated into 1 mL of BHI without antibiotics and incubated for 24 h to generate passage 2 (P2). P2 cultures were stored at −80°C in BHI containing 20% glycerol.

Subsequently, 50 μL of each passage was transferred into 1 mL of fresh antibiotic-free BHI and incubated for 24 h to generate the next passage. This serial passaging was continued through passage 6 (P6), after which cultures were also stored at −80°C in 20% glycerol.

To determine the frequency of kanamycin-resistant cells, frozen stocks of P2 and P6 were inoculated into 1 mL of antibiotic-free BHI in triplicate and incubated for 24 h, generating P3 and P7 cultures, respectively. Cultures were serially diluted in phosphate-buffered saline (PBS), and 2 μL of each dilution was spotted onto BHI agar plates with or without kanamycin (500 μg/mL). Plates were incubated anaerobically at 37°C for 48 h, and CFUs were enumerated. The frequency of kanamycin-resistant cells was calculated as the ratio of CFUs recovered on kanamycin-containing plates to those recovered on antibiotic-free plates.

### Growth assay of mutant strains

To quantify the growth of Δ*immR* and Δ*immR*-*WTCR* strains, passage 2 lineages from the serial-passage experiment described above were cultured overnight in BHI containing kanamycin (500 μg/mL) under anaerobic conditions at 37°C. The resulting overnight cultures (P3) were diluted 1:10 into 150 μL of fresh BHI in 96-well plates containing kanamycin (500 μg/mL). Growth was monitored under aerobic conditions by measuring OD₆₀₀ using an Agilent BioTek Synergy H1 microplate reader at 37°C for 12 h.

For experiments involving translation-associated mutants, cultures were prepared to achieve comparable cell densities before growth assays and spot-dilution analyses because the strains differed substantially in growth rate. Faster-growing strains, including the Δ*fmt* ΔTn*Smu1*, Δ*rnjA* ΔTn*Smu1*, and Δ*rnjB* ΔTn*Smu1* strains, were prepared by inoculating 100 μL of overnight culture into 900 μL of fresh BHI and incubating overnight. Slower-growing Δ*attR* derivatives, including Δ*fmt* Δ*attR*, Δ*rnjA* Δ*attR*, and Δ*rnjB* Δ*attR*, were cultured for 2 days, harvested by centrifugation at 800 × *g* for 1 min, resuspended in 1 mL of fresh BHI, and incubated for an additional 2 days. For growth assays, cultures were inoculated into 150 μL of fresh BHI in 96-well plates and incubated anaerobically at 37°C. WT, ΔTn*Smu1*, and Tn*Smu1*-loss derivatives were inoculated at a 1:1,000 dilution, whereas the slower-growing Δ*attR* derivatives were inoculated at a 1:10 dilution to compensate for their severe growth defects. Viable cell numbers were determined at 0, 24, and 48 h by colony-forming unit (CFU) enumeration.

### Whole-genome sequencing and variant calling

Whole-genome sequencing was performed by SeqCenter (https://www.seqcenter.com/) using 2 × 150-bp paired-end Illumina sequencing. Only genomes with an average sequencing depth of at least 100× were included in downstream analyses.

Raw FASTQ reads were quality trimmed using BBDuk and aligned to the *S. mutans* UA159 reference genome (NC_004350) using Geneious Prime (version 2025.1.2) (https://www.geneious.com/). Structural variants, including copy-number variations and deletions, were identified using the Geneious Prime variant-calling pipeline and manually inspected.

## Data and materials availability

All data needed to evaluate the conclusions in the paper are present in the paper and/or the Supplementary files. The genome sequence data were deposited to BioProject (PRJNA1504854). All bacterial strains generated in this study, including the essential-gene deletion mutants, are available from the authors upon request.

## Acknowledgments

This work was funded by grants from the National Institutes of Health R03DE034511 (L.B.) and R01DE030121 (P.X. and TK). T.K. was also supported by R21DE034103. P.X. was supported by the CCTR Endowment Fund.

---

**Fig. S1.**
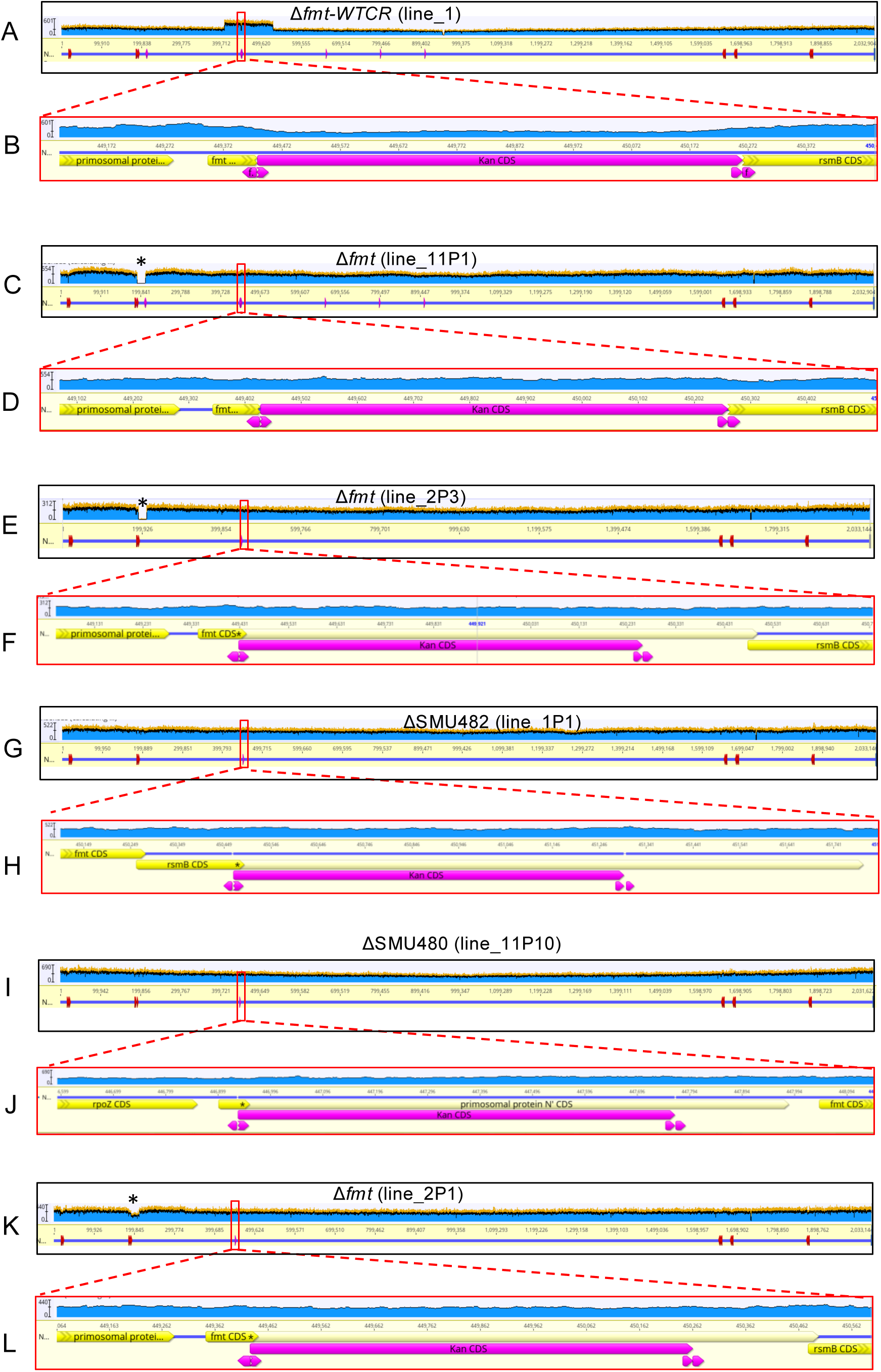
Whole-genome sequencing confirms that loss of *fmt* is associated with Tn*Smu1* destabilization. Sequencing reads were aligned to the *S. mutans* UA159 reference genome (NC_004350), or to the indicated modified reference, to verify the relevant deletion alleles and Tn*Smu1* status. Red boxes in panels A, C, E, and G indicate the regions enlarged in panels B, D, F, and H, respectively. **(A–B)** Whole-genome sequencing of the Δ*fmt*-WTCR strain (line_1), showing retention of a wild-type *fmt* copy. **(C–D)** Whole-genome sequencing of an independently evolved Δ*fmt* lineage (line_11P1), confirming the Δ*fmt* allele and showing reduced Tn*Smu1* coverage at passage 1. **(E–F)** Whole-genome sequencing of Δ*fmt* lineage line_2 at passage 3 (line_2P3), showing further reduction/loss of Tn*Smu1* coverage. **(G–H)** Whole-genome sequencing of the ΔSMU482 mutant (line_1P1), confirming deletion of the downstream SMU482 gene without destabilization of Tn*Smu1*. **(I–J)** Whole-genome sequencing of the ΔSMU480 mutant (line_11P10), confirming deletion of the upstream SMU480 gene without destabilization of Tn*Smu1*. **(K–L)** Whole-genome sequencing of the same Δ*fmt* lineage shown in E–F at passage 1 (line_2P1), showing a partial reduction in Tn*Smu1* coverage before the more pronounced loss observed at passage 3.

**Fig. S2.**
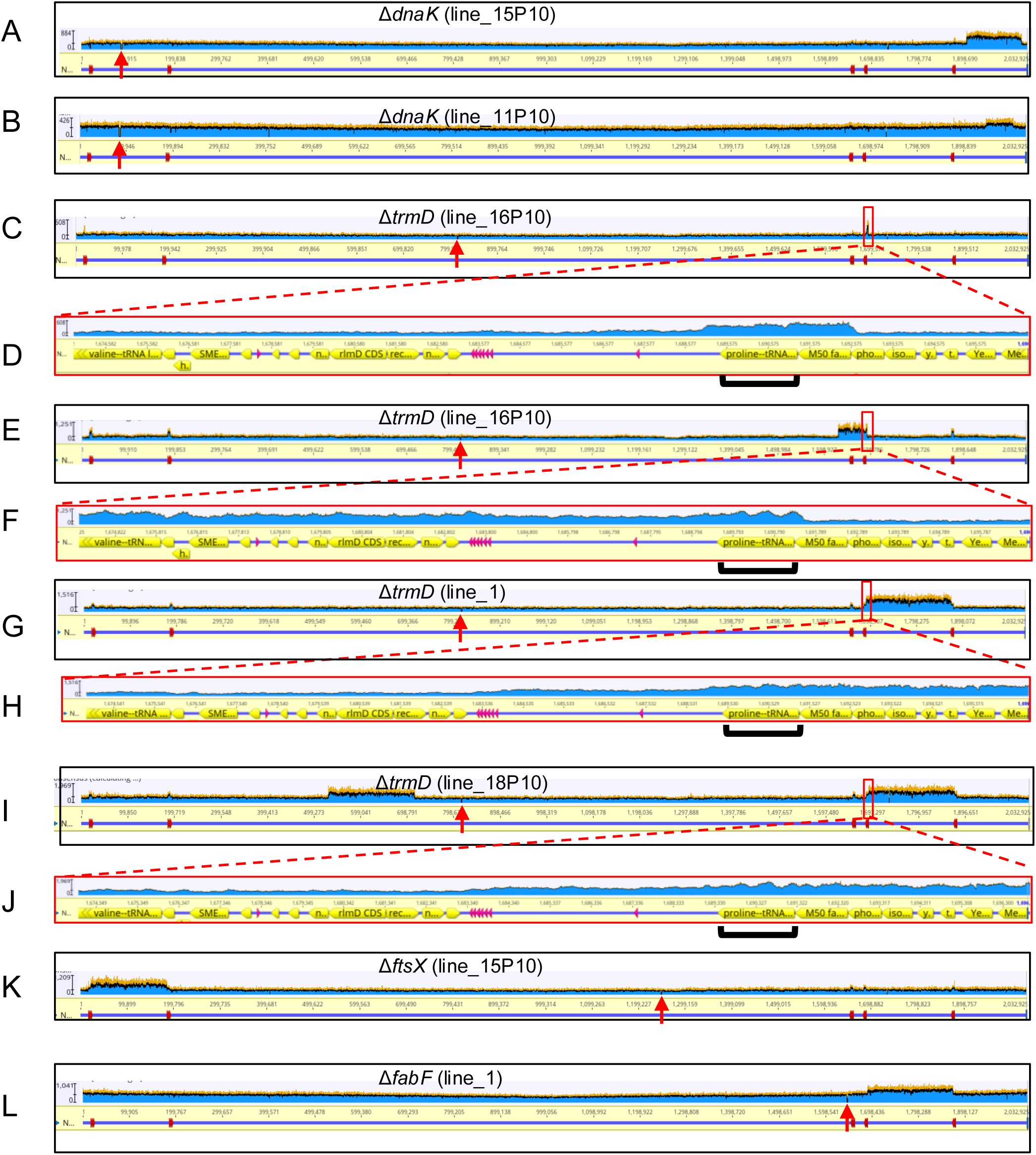
Whole-genome sequencing of evolved control lineages demonstrates adaptive genome evolution without Tn*Smu1* loss. Whole-genome sequencing was performed on independently evolved control mutants affecting protein folding, tRNA modification, cell division, or fatty acid biosynthesis. **(A–B)** Whole-genome sequencing coverage profiles of two independently evolved Δ*dnaK* lineages. **(C–J)** Whole-genome sequencing coverage profiles of four independently evolved Δ*trmD* lineages. Red boxes in panels C, E, G, and I indicate the genomic regions enlarged in panels D, F, H, and J, respectively, highlighting representative chromosomal copy-number amplifications. **(K)** Whole-genome sequencing coverage profile of an evolved Δ*ftsX* lineage. **(L)** Whole-genome sequencing coverage profile of an evolved Δ*fabF* lineage. None of these evolved lineages exhibited loss or amplification of the Tn*Smu1* region. Despite the accumulation of adaptive genomic changes, including recurrent chromosomal copy-number amplifications of these lineages, Tn*Smu1* remained intact in all lineages shown. Red arrows indicate the locations of the intended gene deletions. The height of the blue coverage profile indicates the number of sequencing reads mapped to the reference genome at the coordinates shown on the x axis. A doubling of sequence reads indicates duplication of the affected region.

**Fig. S3.**
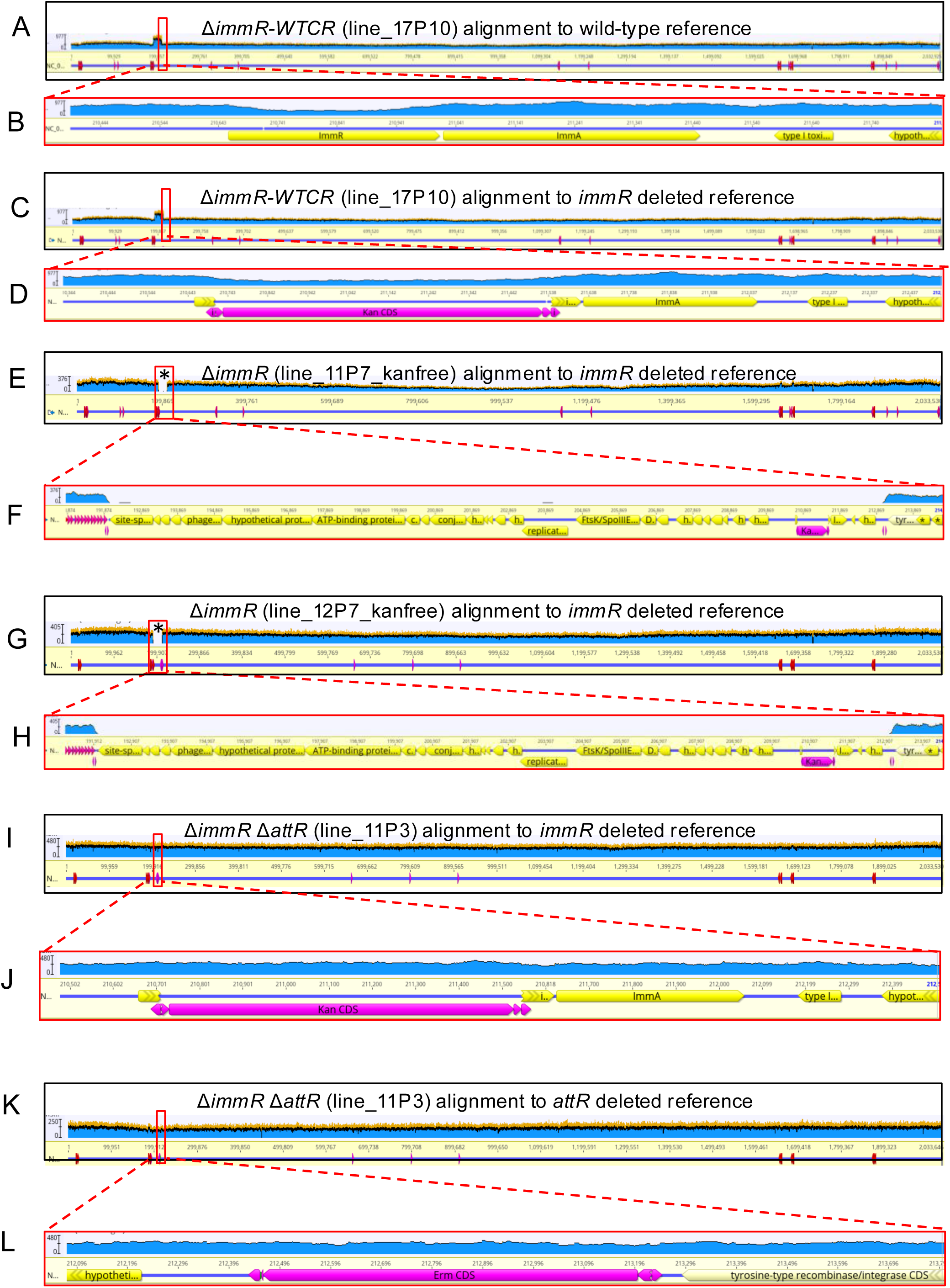
Whole-genome sequencing confirms that *immR* deletion drives Tn*Smu1* loss, whereas deletion of *attR* prevents Tn*Smu1* loss. Sequencing reads were aligned to modified *S. mutans* UA159 reference genomes (NC_004350), as indicated for each panel. Red boxes in panels A, C, E, G, I, and K indicate the regions enlarged in panels B, D, F, H, J, and L, respectively. **(A–B)** Whole-genome sequencing of the Δ*immR*-WTCR strain (line_17P10) aligned to the wild-type reference genome, confirming retention of a wild-type *immR* allele. **(C–D)** Whole-genome sequencing of the same Δ*immR*-WTCR strain (line_17P10) aligned to the modified reference genome containing the Δ*immR*::*kan* allele, confirming the presence of the Δ*immR*::*kan* allele. Together, A–D demonstrate that the Δ*immR*-WTCR strain retains both a wild-type *immR* copy and the Δ*immR*::*kan* allele. **(E–F)** Whole-genome sequencing of the kanamycin-sensitive isolate line_11P7_kanfree, recovered from an independently evolved Δ*immR* lineage and aligned to the Δ*immR* reference genome, confirming loss of the Tn*Smu1* region. **(G–H)** Whole-genome sequencing of the kanamycin-sensitive isolate line_12P7_kanfree, independently derived from another evolved Δ*immR* lineage and aligned to the Δ*immR* reference genome, likewise confirming loss of Tn*Smu1*. Two additional independently evolved Δ*immR* lineages (line _15P7_kanfree and line _16P7_kanfree) with Tn*Smu1* loss are not shown. **(I–J)** Whole-genome sequencing of the passaged Δ*immR* Δ*attR* strain (line_11P3) aligned to the modified reference genome containing the Δ*immR*::*kan* allele, confirming retention of the Δ*immR*::*kan* allele and Tn*Smu1*. **(K–L)** Whole-genome sequencing of the same Δ*immR* Δ*attR* strain (line_11P3) aligned to the modified reference genome containing the Δ*attR*::*erm* allele, confirming retention of the expected Δ*attR* allele.

**Fig. S4.**
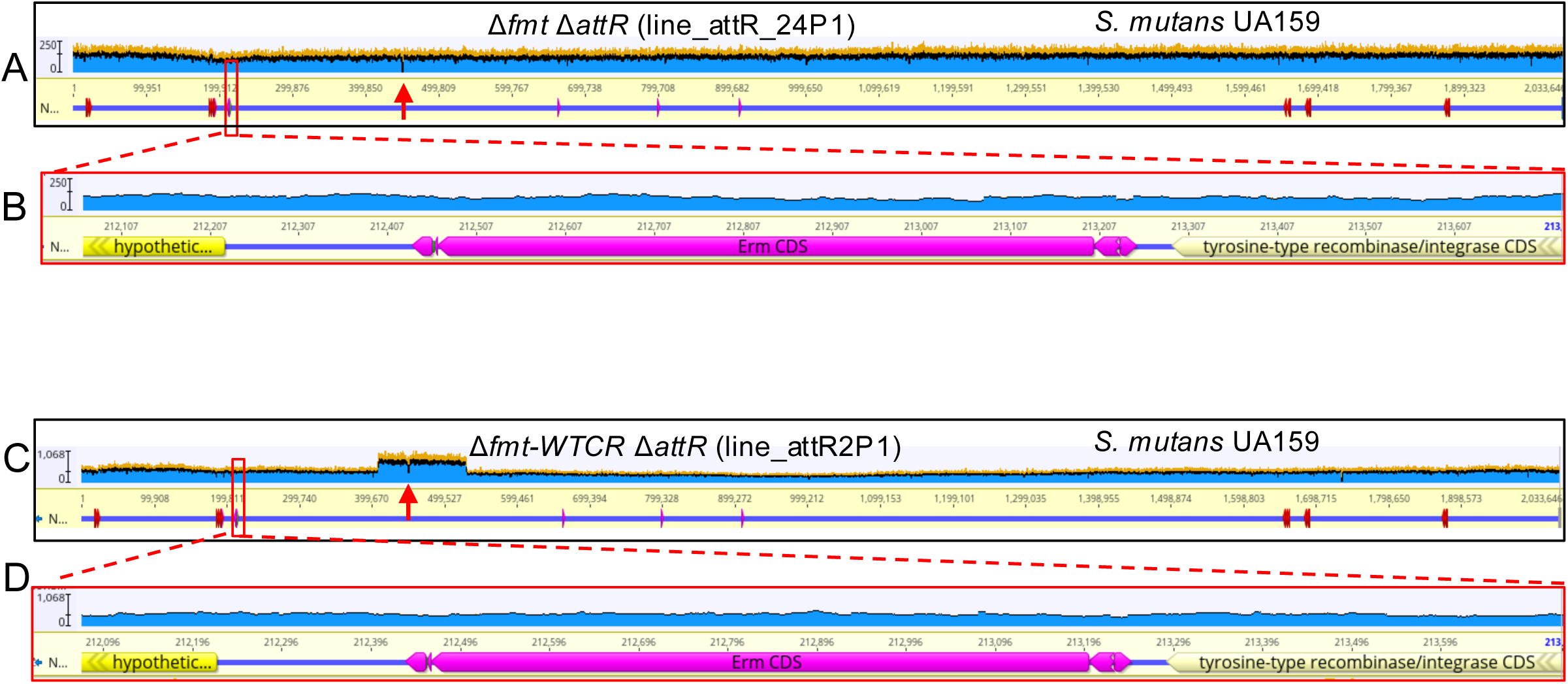
Whole-genome sequencing confirms that *fmt* deletion does not induce Tn*Smu1* loss in the absence of *attR*. Sequencing reads were aligned to a modified *S. mutans* UA159 reference genome (NC_004350) in which the *attR* site was replaced with an erythromycin-resistance (*erm*) cassette. Red boxes in panels **A** and **C** indicate the *attR* replacement region enlarged in panels **B** and **D**, respectively. (A–B) Whole-genome sequencing of the passaged Δ*fmt* Δ*attR* strain. (C–D) Whole-genome sequencing of the passaged Δ*fmt*-*WTCR* Δ*attR* strain.

**Fig. S5.**
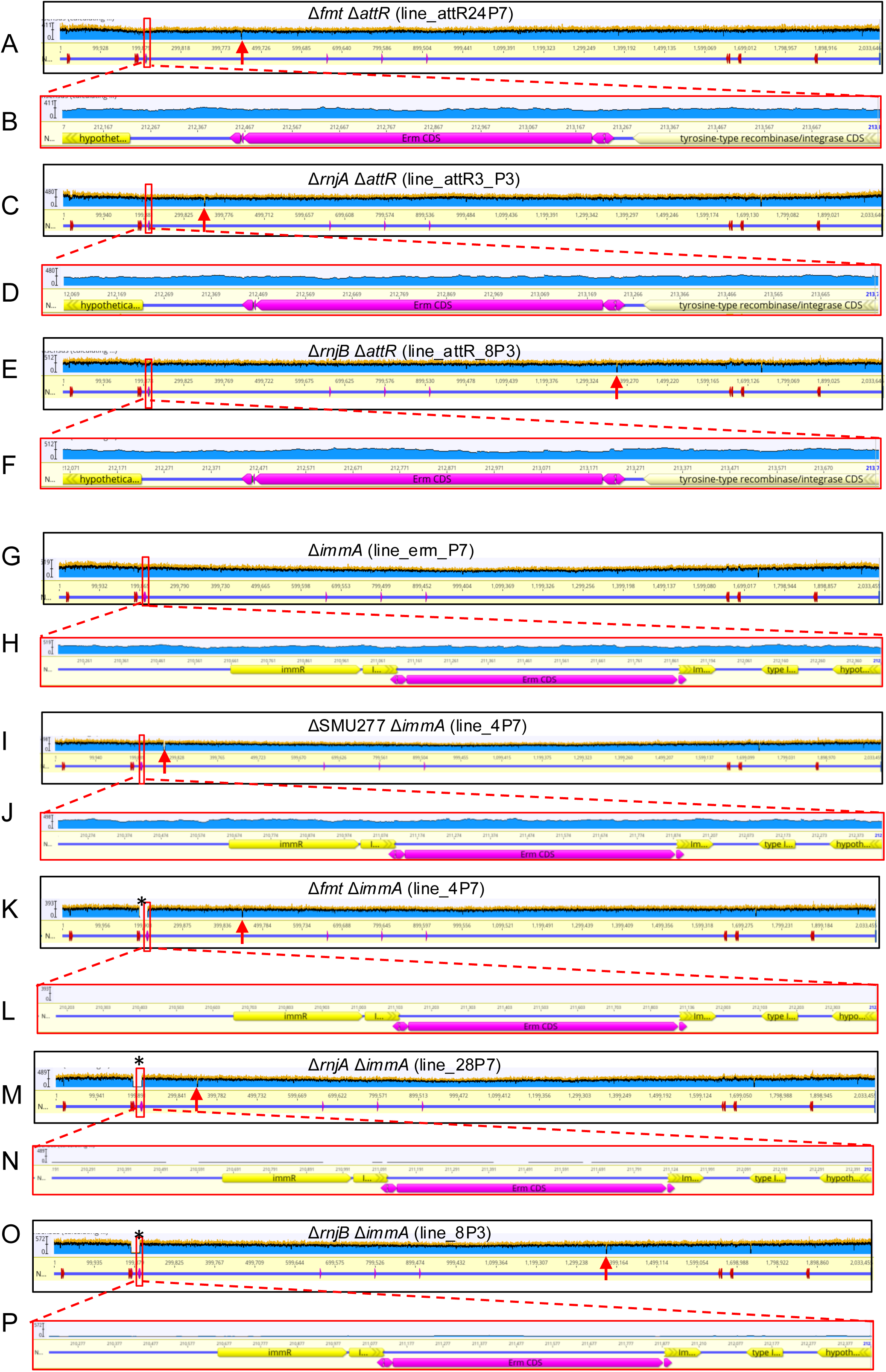
Whole-genome sequencing confirms Tn*Smu1* status in passaged Δ*attR* and Δ*immA* mutants following disruption of *fmt, rnjA,* and *rnjB*. Sequencing reads were aligned to modified *S. mutans* UA159 reference genomes (NC_004350), in which either the *attR* site (**A–F**) or the *immA* coding sequence (G–P) was replaced with an erythromycin-resistance (*erm*) cassette. Red boxes in panels **A, C, E, G, I, K, M,** and **O** indicate the regions enlarged in panels **B, D, F, H, J, L, N**, and **P**, respectively. **(A–F)** Whole-genome sequencing of passaged Δ*attR*/translation-associated double mutants: Δ*fmt* Δ*attR* (**A–B**), Δ*rnjA* Δ*attR* (**C–D**), and Δ*rnjB* Δ*attR* (**E–F**). **(G–P)** Whole-genome sequencing of passaged Δ*immA* mutants: Δ*immA* (**G, H**), Δ*SMU277* Δ*immA* (**I–J**), Δ*fmt* Δ*immA* (**K–L**), Δ*rnjA* Δ*immA* (**M–N**), and Δ*rnjB* Δ*immA* (**O–P**). Loss of the Tn*Smu1* region is indicated by asterisks in panels K, M, and O, whereas the Tn*Smu1* locus remains intact in the other strains. Red arrows indicate the intended deletion regions. Vertical red arrow, site of original gene deletion; the height of the blue segments indicates the number of sequence reads mapped to the reference sequence at the coordinates shown on the X axis. Regions marked by asterisks indicate loss of the ∼20-kb Tn*Smu1* element.

**Fig. S6.**
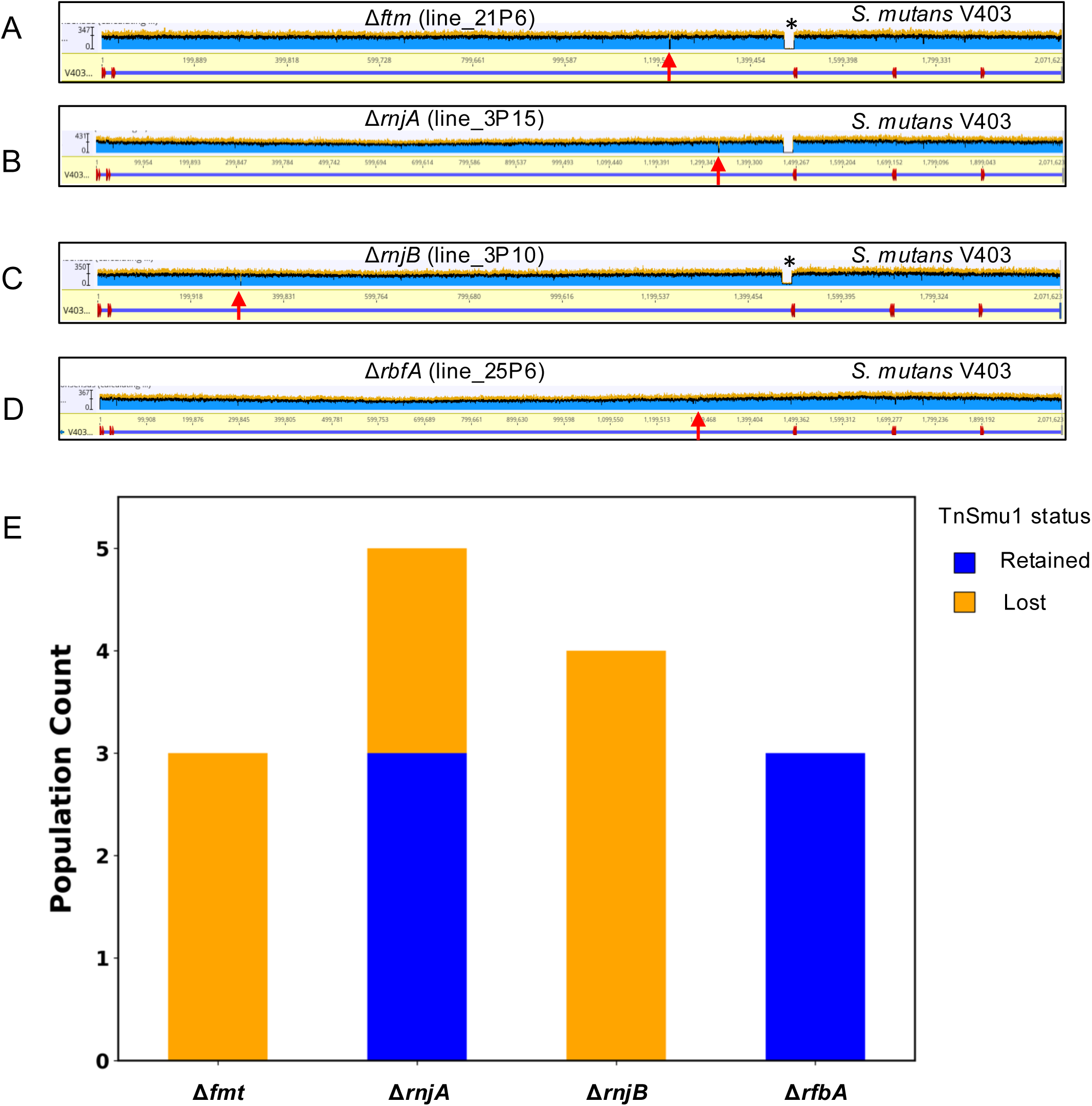
Host translation influences Tn*Smu1* stability across genetically distinct *S. mutans* strains. **(A-D),** Whole-genome sequencing confirming Tn*Smu1* status after deletion of *fmt* (**A**), *rnjA* (**B**), *rnjB* (**C**) and *rbfA* (**D**) in strain V403. Red arrows indicate the intended deletion regions. Vertical red arrow, site of original gene deletion; the height of the blue segments indicates the number of sequence reads mapped to the reference sequence at the coordinates shown on the X axis. Regions marked by asterisks indicate loss of the ∼20-kb Tn*Smu1* element. **(E),** Tn*Smu1* loss frequencies in the clinical isolate V403 following disruption of *fmt*, *rnjA*, *rnjB* and *rbfA*.

## Supplemental Tables

Table S1: Whole-genome sequencing analysis of Tn*Smu1* stability in evolved deletion lineages in *S. mutans* UA159.

Table S2. Quantification of Tn*Smu1* loss in Δ*immR* lineages over serial passage (P3 vs. P7) via kanamycin resistance retention profiling.

Table S3. Quantitative growth evaluation of Δ*immR* in *S. mutans* UA159 strains.

Table S4. Quantitative fitness evaluation and growth recovery of *S. mutans* UA159 strains upon Tn*Smu1* loss during *immR* perturbation.

Table S5. Quantitative fitness evaluation and growth recovery of *S. mutans* UA159 strains upon Tn*Smu1* loss during *fmt* perturbation.

Table S6. Quantitative fitness evaluation and growth recovery of *S. mutans* UA159 strains upon Tn*Smu1* loss during *rnjA* or *rnjB* perturbation.

Table S7. Whole-genome sequencing analysis of Tn*Smu1* stability following disruption of translation-associated host genes in the Δ*attR* background.

Table S8. Whole-genome sequencing analysis of Tn*Smu1* stability following disruption of translation-associated host genes in the Δ*immA* background.

Table S9. Whole-genome sequencing analysis of Tn*Smu1* stability in translation-associated mutants of *S. mutans* V403.

Table S10: Primers used in this study.

## References

1. Tokuda M, Shintani M. 2024. Microbial evolution through horizontal gene transfer by mobile genetic elements. Microb Biotechnol 17:e14408.

2. Gomberg AF, Grossman AD. 2024. It’s complicated: relationships between integrative and conjugative elements and their bacterial hosts. Curr Opin Microbiol 82:102556.

3. Burch CL, Romanchuk A, Kelly M, Wu Y, Jones CD. 2023. Empirical Evidence That Complexity Limits Horizontal Gene Transfer. Genome Biol Evol 15.

4. Arnold BJ, Huang IT, Hanage WP. 2022. Horizontal gene transfer and adaptive evolution in bacteria. Nat Rev Microbiol 20:206–218.

5. Partridge SR, Kwong SM, Firth N, Jensen SO. 2018. Mobile Genetic Elements Associated with Antimicrobial Resistance. Clin Microbiol Rev 31.

6. Weisberg AJ, Chang JH. 2023. Mobile Genetic Element Flexibility as an Underlying Principle to Bacterial Evolution. Annu Rev Microbiol 77:603–624.

7. Delavat F, Miyazaki R, Carraro N, Pradervand N, van der Meer JR. 2017. The hidden life of integrative and conjugative elements. FEMS Microbiol Rev 41:512–537.

8. Cury J, Touchon M, Rocha EPC. 2017. Integrative and conjugative elements and their hosts: composition, distribution and organization. Nucleic Acids Res 45:8943–8956.

9. McLellan LK, Anderson ME, Grossman AD. 2022. TnSmu1 is a functional integrative and conjugative element in Streptococcus mutans that when expressed causes growth arrest of host bacteria. Mol Microbiol 118:652–669.

10. King S, Quick A, King K, Walker AR, Shields RC. 2022. Activation of TnSmu1, an integrative and conjugative element, by an ImmR-like transcriptional regulator in *Streptococcus mutan*s. Microbiology (Reading) 168:001254.

11. Garriss G, Waldor MK, Burrus V. 2009. Mobile Antibiotic Resistance Encoding Elements Promote Their Own Diversity. PLoS Genet. 5(12): e1000775.

12. McKeithen-Mead S, Anderson ME, García-Heredia A, Grossman AD. 2025. Activation and modulation of the host response to DNA damage by an integrative and conjugative element. J Bacteriol 207:e0046224.

13. Duval M, Simonetti A, Caldelari I, Marzi S. 2015. Multiple ways to regulate translation initiation in bacteria: Mechanisms, regulatory circuits, dynamics. Biochimie 114:18–29.

14. Starosta AL, Lassak J, Jung K, Wilson DN. 2014. The bacterial translation stress response. FEMS Microbiol Rev 38:1172–201.

15. Webster MW. 2025. Initiation of Translation in Bacteria and Chloroplasts. J Mol Biol 437:169137.

16. Ajdić D, McShan WM, McLaughlin RE, Savić G, Chang J, Carson MB, Primeaux C, Tian R, Kenton S, Jia H, Lin S, Qian Y, Li S, Zhu H, Najar F, Lai H, White J, Roe BA, Ferretti JJ. 2002. Genome sequence of Streptococcus mutans UA159, a cariogenic dental pathogen. Proc Natl Acad Sci U S A 99:14434–9.

17. Bao L, Zhu Z, Ismail A, Zhu B, Anandan V, Whiteley M, Kitten T, Xu P. 2025. Experimental evolution of gene essentiality in bacteria. mBio 16:e0300525.

18. Shields RC, Zeng L, Culp DJ, Burne RA. 2018. Genomewide Identification of Essential Genes and Fitness Determinants of *Streptococcus mutans* UA159. mSphere 3.

19. Menard KL, Grossman AD. 2013. Selective pressures to maintain attachment site specificity of integrative and conjugative elements. PLoS Genet 9:e1003623.

20. Lahry K, Datta M, Varshney U. 2024. Genetic analysis of translation initiation in bacteria: An initiator tRNA-centric view. Mol Microbiol 122:772–788.

21. Chen X, Liu N, Khajotia S, Qi F, Merritt J. 2015. RNases J1 and J2 are critical pleiotropic regulators in Streptococcus mutans. Microbiology (Reading) 161:797–806.

22. Durand S, Gilet L, Bessières P, Nicolas P, Condon C. 2012. Three essential ribonucleases-RNase Y, J1, and III-control the abundance of a majority of Bacillus subtilis mRNAs. PLoS Genet 8:e1002520.

23. Sharma IM, Woodson SA. 2020. RbfA and IF3 couple ribosome biogenesis and translation initiation to increase stress tolerance. Nucleic Acids Res 48:359–372.

24. Clifton BE, Fariz MA, Uechi G-I, Laurino P. 2021. Evolutionary repair reveals an unexpected role of the tRNA modification m1G37 in aminoacylation. Nucleic Acids Res 49:12467–12485.

25. Bao L, Bradley JL, Anandan V, Tyc KM, Zhu Z, Vossen JA, Assi VF, Benbei JS, Zollar N, Kitten T, Xu P. 2026. A genome-wide in vivo screen reveals fitness pathways required for streptococcal infective endocarditis. PLoS Pathog 22:e1014156.

26. Ramsay JP, Ronson CW. 2015. Silencing quorum sensing and ICE mobility through antiactivation and ribosomal frameshifting. Mob Genet Elements 5:103–108.

27. Paik S, Brown A, Munro CL, Cornelissen CN, Kitten T. 2003. The sloABCR operon of Streptococcus mutans encodes an Mn and Fe transport system required for endocarditis virulence and its Mn-dependent repressor. J Bacteriol 185:5967–75.

28. Bose B, Auchtung JM, Lee CA, Grossman AD. 2008. A conserved anti-repressor controls horizontal gene transfer by proteolysis. Mol Microbiol 70:570–582.

29. Carraro N, Libante V, Morel C, Decaris B, Charron-Bourgoin F, Leblond P, Guédon G. 2011. Differential regulation of two closely related integrative and conjugative elements from *Streptococcus thermophilus*. BMC Microbiol 11:238.

30. Haag AF, Podkowik M, Ibarra-Chávez R, Gallego del Sol F, Ram G, Chen J, Marina A, Novick RP, Penadés JR, et al. 2021. A regulatory cascade controls Staphylococcus aureus pathogenicity island activation. Nat Microbiol 6:1300–1308.

31. Tang L, Yang W, Yang L, Lv Y, Zhang J. 2026. Targeting Horizontal Gene Transfer to Combat Antimicrobial Resistance: A Review of Mechanisms, Drivers, and Multi-Omics Strategies. Infect Drug Resist 19:589962.

32. Kitten T, Michalek LC, Munro MS, Francis LM. 2000. Genetic characterization of a Streptococcus mutans LraI family operon and role in virulence. Infection and Immunity 68:4441.

33. Xu P, Ge X, Chen L, Wang X, Dou Y, Xu JZ, Patel JR, Stone V, Trinh M, Evans K, Kitten T, Bonchev D, Buck GA. 2011. Genome-wide essential gene identification in *Streptococcus sanguinis*. Sci Rep 1:125.

